# The effect of deferring feedback on rule-based and information-integration category learning

**DOI:** 10.1101/2024.10.31.621443

**Authors:** C. E. R. Edmunds, Kathryn Carpenter, Andy J. Wills, Fraser Milton

## Abstract

Previous work has shown that deferring feedback significantly impairs two-dimensional information-integration category learning, often thought to recruit an implicit learning system, but leaves intact unidimensional rule-based learning, commonly assumed to engage an explicit system (Smith et al., 2014). These results were taken to support the influential COmpetition between Verbal and Implicit Systems (COVIS) dual-process theory. This conclusion has subsequently been challenged by the finding that this dissociation disappears when the number of relevant dimensions is matched between tasks (Le Pelley et al., 2019). However, as well as replacing a unidimensional rule-based task with a two-dimensional conjunction task, Le Pelley et al. also changed the stimuli that were used in their study making it unclear which of these alterations was driving the difference in results. The current paper directly examined how both category structure and stimulus type influence the deferred feedback effect. We replicated both the original sets of results but found that deferred feedback also impaired information-integration learning to a greater extent than a conjunction task when Smith et al.’s original stimuli were used. This result suggests that the effect of deferred feedback on category learning is more complicated than has previously been documented and highlights the critical role the choice of stimuli has in determining whether the effect is obtained.

---

Category learning is a fundamental cognitive skill that enables us to function effectively in our everyday environment. Classifying items into meaningful groups enables us to react to the virtually infinite amount of novel items we come across in our daily lives in an appropriate manner rather than having to learn about every item individually. The importance of categorization to human functioning is reflected in the considerable amount of attention that has been paid in recent decades to the processes that underlie category learning. Perhaps the most enduring and contentious debate in the field is the question of whether category learning is best explained by dual process theories that propose categorization can be the result of both explicit and implicit learning systems (e.g., Ashby, Alfonso-Reese, Turken, & Waldron, 1998; Ashby & Maddox, 2011) or single system accounts that argue that categorization can most parsimoniously be explained as the result of a single explicit system (e.g., Newell, Dunn, & Kalish, 2011, Wills, et al., 2019).

The COVIS (COmpetition between Verbal and Implicit Systems) model (Ashby et al., 1998) is arguably the most influential dual process account of category learning. COVIS assumes that people have two parallel systems of category learning. The explicit system is rule-based and requires working memory to generate and test rules. The explicit system is consequently thought to be recruited for easy to verbalise rule-based (RB) category structures such as unidimensional (UD) or conjunction (CJ) rules, shown in Figures 1a and 1b. The implicit system combines information from multiple dimensions predecisionally through immediate feedback to create stimulus-response associations. The implicit system is therefore thought to be preferentially engaged when category structures are difficult or impossible to verbalize because the explicit system is not effective at learning these types of structures. COVIS research has typically used an information-integration (II) task as shown in Figure 1c to encourage use of the implicit system because the optimal diagonal decision boundary cannot easily be captured by a rule.

**Fig 1.**
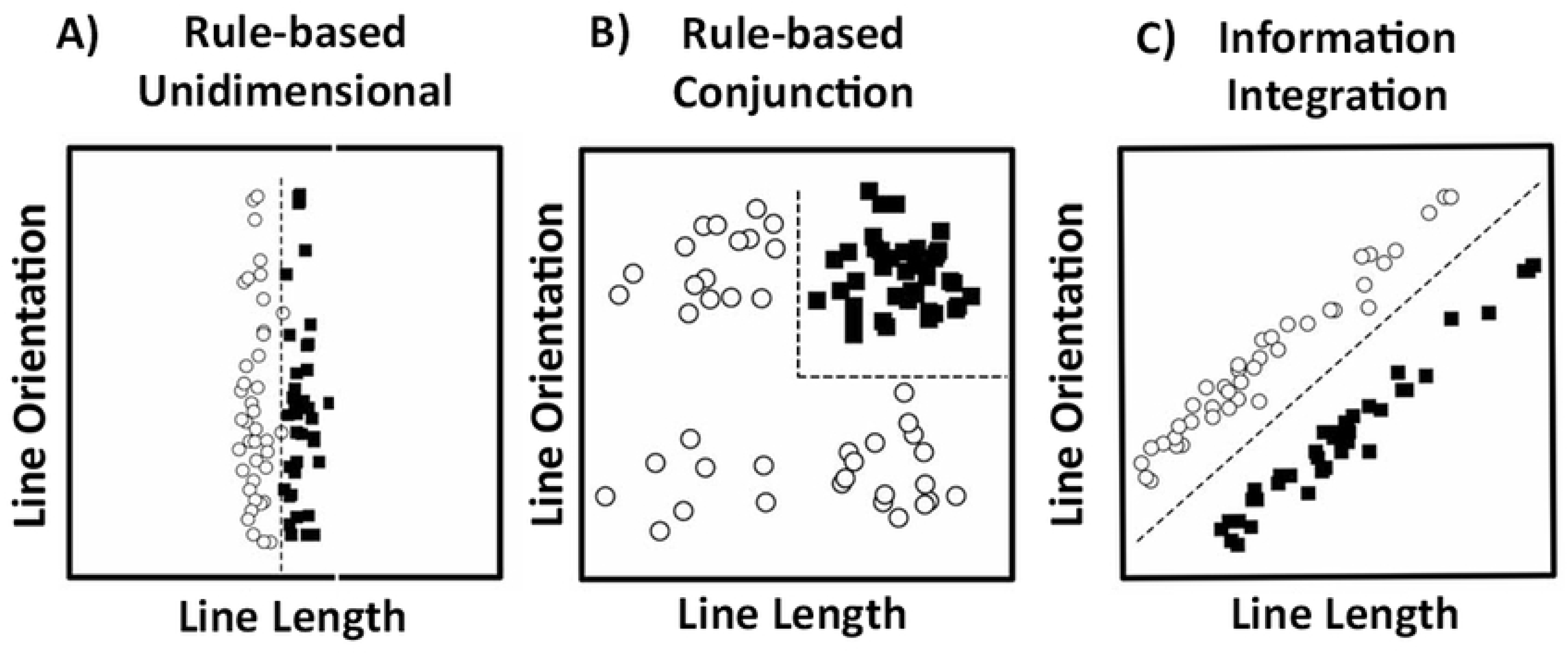
Examples of unidimensional, conjunction, and information-integration category structures. Each open circle represents one member of category A; each filled square represents one member of category B. Figure adapted from Wills et al. (2013) and Zeithamova and Maddox (2006).

These category structures have revealed a wide range of behavioral dissociations, which have been interpreted as evidence supporting COVIS (for reviews, see Ashby & Maddox, 2011; Smith & Church, 2018). However, a number of these dissociations have subsequently been criticised either for being driven by methodological flaws (e.g., Newell, Moore, Wills & Milton, 2013) or for failing to replicate (e.g. Edmunds, Milton, & Wills, 2019) which means that this debate is still far from settled.

The current work therefore aims to extend this previous research by evaluating a particularly intriguing behavioral dissociation. Smith et al. (2014) looked at the effect of different forms of feedback on learning two different category structures. They presented participants with either a UD RB or an II category structure. Participants either received immediate feedback after every trial, or deferred feedback collectively presented after every six categorization trials. There was a significant interaction - II participants receiving deferred feedback were impaired compared to II participants who received immediate feedback, yet RB participants were unaffected by the feedback manipulation. Smith et al. argue that this dissociation provides some of the strongest evidence to date for the existence of separate explicit and implicit category learning systems.

COVIS explains this pattern of results by proposing that RB learning should remain intact when receiving deferred feedback as participants would optimally engage the explicit system which stores the responses made to each trial, as well as the category learning rule, in working memory. This enables the explicit system to hold category learning information for later use, and it is therefore unimpaired by deferred feedback as information can be stored until the later feedback presentation. This assumption led the authors to predict that RB learning would ‘flourish’ (Smith et al., 2014, p. 450) in deferred feedback conditions. However, COVIS also predicts that II learning would be disadvantaged by deferred feedback as optimal II learning is performed by the implicit system, which procedurally associates the representations of the perceived stimulus and the response made through a dopamine release initiated when presented with positive feedback. However, if feedback is delayed or deferred then the neural representation of the response made and the stimulus seen will have decayed, and therefore the representations cannot be associated when dopamine is released (Ashby et al., 1998).

There has, however, been extensive debate on the merits of comparing an II task to a UD RB task. Dual-process proponents argue that these category structures act as “elegant mutual controls” (Smith et al., 2014, p. 249) as they are simple rotations of one another, matched in terms of category size and number of categories, as well as within-category similarity and between-category separation. On the other hand, critics point out that these structures are well matched on everything *apart from the number of dimensions relevant for categorising*. In the UD structure, only one dimension is relevant, whilst in the II structure both dimensions are relevant leading to the claim that the II task is more cognitively difficult than the UD task. This is consistent with the common finding that the II condition typically leads to poorer performance than the UD condition (e.g., Ashby, Ell, & Waldron, 2003; Ashby, Maddox, & Bohil, 2002) which could potentially be the cause of dissociations found where this is the case. According to this view, a two-dimensional CJ RB structure (as in Figure 1b) provides a more suitable comparison to the two-dimensional II structure than a UD RB task. We note that COVIS proponents also believe that a CJ structure taps into the explicit system and have used it in their own research (e.g., Filoteo, Lauritzen, & Maddox, 2010; Helie & Ashby, 2012; Zeithamova & Maddox, 2006). Previous work has shown that when a CJ structure is used instead of a UD structure, dissociations between RB and II category learning disappear (e.g., Carpenter et al., 2016; Edmunds, Milton, & Wills, 2015).

This critique motivated Le Pelley, Newell, and Nosofsky (2019) to provide a first evaluation of Smith et al.’s (2014) experiment. In a between-subjects study, they compared the effect of immediate and deferred feedback on the learning of UD, CJ and II category structures. They replicated the original dissociation between the UD and II category structure. However, they also found a significant interaction between feedback type and the UD and CJ category structures: the negative impact of deferred feedback was significantly greater in the CJ condition than the UD condition. In contrast, there was no difference in the negative effect of deferred feedback between the CJ and II conditions.

This is consistent with the idea that the original dissociation was driven by a difficulty effect. Deferred feedback disrupted the II condition, which they argued requires high memory demands to combine multiple dimensions and remember into which category such a combination has previously been assigned. In contrast the UD condition involves lower memory demands to store the optimal one-dimensional rule. These results appear to pose a challenge to the dual process account proposed by COVIS.

However, there are grounds to believe that such a conclusion may be premature. One notable aspect of both the Smith et al. (2014) and Le Pelley et al., (2019) studies is that they each relied on a single unreplicated study. Given that in recent times there has been increased recognition of the importance of replicating results to ensure their robustness this seems a limitation in both studies. Whilst it is true that Le Pelley et al. provided a conceptual replication of Smith et al.’s dissociation between the UD and II category structures, they used a different type of stimuli. Specifically, Smith et al. used stimulus rectangles varying in size and dot density (see Figure 2) – which we call the “dot stimuli”. In contrast, Le Pelley et al. used simpler “line stimuli” varying in length and orientation. Whilst the stimuli Le Pelley et al. used have frequently been used in previous COVIS related research (e.g., Filoteo, Lauritzen, & Maddox, 2010; Filoteo, Maddox, Salmon, & Song, 2005; Maddox et al., 2010) they are extremely different from the more perceptually complex dot stimuli used by Smith et al.. This means that there is a second potentially important difference between the studies outside of the introduction of the CJ category structure which potentially could be contributing to the difference in findings and which consequently arguably should have been more tightly controlled.

**Fig 2.**
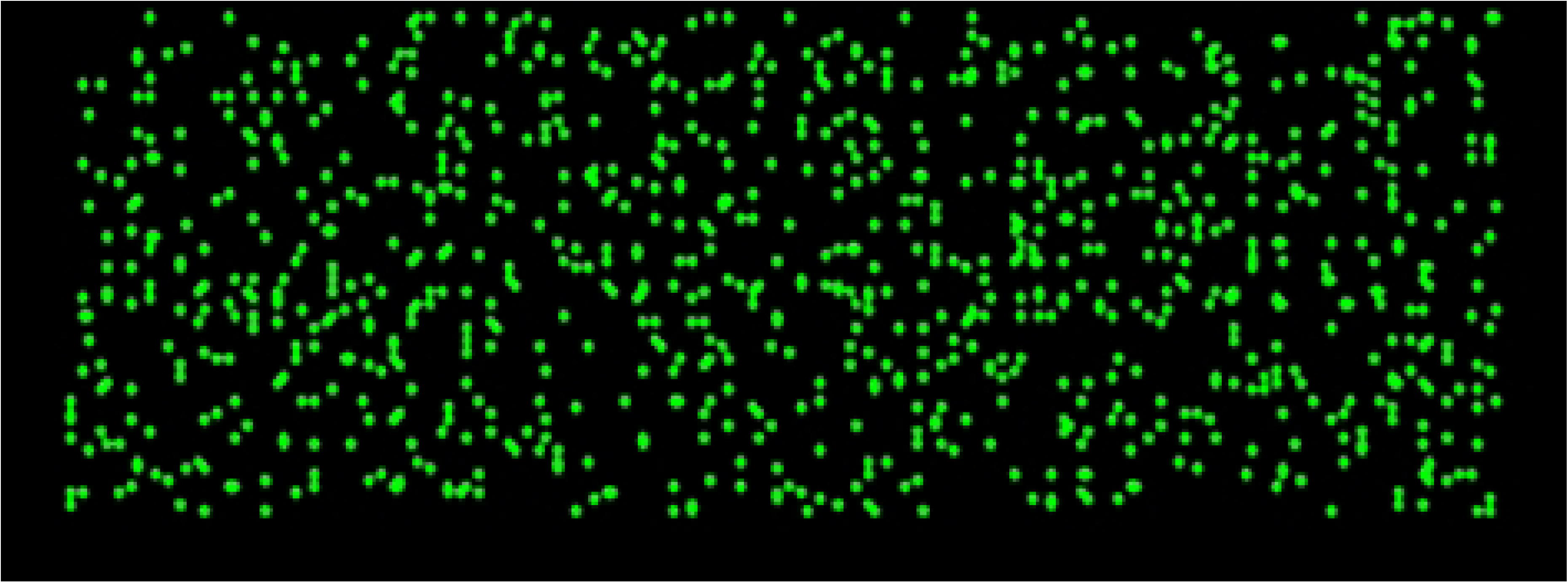
Example of the stimuli used in Smith et al. (2014) and Experiments 1 and 3.

This potential issue is underscored by the differing pattern of results for the II and UD conditions in the Smith et al. (2014) and Le Pelley et al. (2019) papers. Le Pelley et al. found that in the immediate feedback control conditions, overall accuracy was lower for the II structure than the UD structure. This is, as discussed above, consistent with past work showing that participants typically find the II structure more difficult to learn than the UD structure. However, Smith et al. found no significant difference in accuracy between II and UD learning under immediate feedback conditions. Their observing a greater effect of deferred feedback for II than UD category learning consequently appears contrary to the difficulty-based explanation outlined earlier. Le Pelley et al. plausibly argued that this difference may reflect that the greater number of participants they recruited in their study gave them greater power to detect subtle effects than Smith et al., but nonetheless it is also suggestive that the differing nature of the stimuli may have an influence on the results and this warrants further investigation. In summary, the question we hope to answer is, do the results of Le Pelley et al. and Smith et al. differ because they used different category structures, or because they used different physical instantiations of those structures? While the latter possibility would still speak to the generalizability of Smith et al.’s findings across stimulus types, the former possibility would support Le Pelley et al.’s stronger claim that Smith et al.’s results are an artefactual side effect of failing to match for the number of relevant stimulus dimensions.

## Experiment 1

In Experiment 1, we directly replicated Smith et al.’s (2014) experiment with participants learning either an II or UD category structure under either deferred or immediate feedback. The category structures were generated by sampling from two multivariate normal distributions, one for each category, rather than the category structures used by Le Pelley et al. (2019) which sampled from two multivariate normal distributions for each category. Note this difference in structure is unlikely to make a difference; however, in the efforts to run an *exact* replication we tried to match Smith et al. as closely as possible. More importantly we used Smith et al.’s dot stimuli varying in size of rectangle and pixel density rather than the lines varying in length and orientation as in Le Pelley et al.

## Method

### Participants and Design

87 University of Exeter students completed the experiment (19 males, mean age 20.24 years, SD=5.32). This sample size was chosen as being similar to that used in Smith et al.’s (2014) study. One participant was excluded because they failed to complete the experiment. Participants received course credit or £5 for their participation. This and Experiments 2a, 2b and 3a received ethical approval from the University of Exeter Psychology Ethics Committee.

The experiment had a 2 (Category structure: UD, II) x 2 (Feedback: Immediate, Deferred) between-subjects design. Participants were randomly assigned to each of the four conditions: UD-immediate (*N* = 20), UD-deferred (*N* = 21), II-immediate (*N* = 20), and II- deferred (*N* = 24).

### Stimuli

The stimuli were the ‘dot stimuli’ used by Smith et al. (2014, see Figure 2). They were unframed rectangles of green dots displayed in a black background that varied in the size of the rectangle and the density of the green dots within the rectangle. In the UD conditions, the size of the rectangle was the relevant dimension for categorisation, but for the II conditions both dimensions were relevant. Following Smith et al., 600 stimuli were generated individually for each participant. Thus, whilst participants in each condition had stimuli generated from the same distributions (see Table 1), the precise visual stimuli presented varied between each individual. The values for the stimuli for each category were drawn from a bivariate normal distribution and rounded to the nearest whole number.

**Table 1.** Distributional characteristics of the unidimensional RB and II category structures in Experiment 1.

| Task | Category | MeanX | MeanY | VarX | VarY | CovXY |
| --- | --- | --- | --- | --- | --- | --- |
| UD | A | 35.86 | 50 | 16.33 | 355.55 | 0 |
|  | B | 64.14 | 50 | 16.33 | 355.55 | 0 |
| II | A | 40.00 | 60.00 | 185.94 | 185.94 | 169.61 |
|  | B | 60.00 | 40.00 | 185.94 | 185.94 | 169.61 |

Note. The numbers define the bivariate normal distribution from which stimuli were sampled. UD = Unidimensional, II = Information-integration, X = rectangle size, Y = dot density, Var = Variance, Cov = Covariance. The values of the stimulus dimensions are in arbitrary units. As in Smith et al. (2014), both dimensions had 101 levels (Levels 0–100). Rectangle width and height (in screen pixels) were calculated as $100 + \text{level}$ and $50 + \text{level}/2$ , respectively. Pixel density was calculated as $0.05 \times 1.018^{\text{level}}$ .

### Procedure

Participants were informed that on each trial they would see a novel stimulus which could be categorized as either A or B, and they were asked to learn into which of two categories a series of stimuli belonged through trial and error. Every trial began with a white fixation cross displayed for 500ms in the center of a black screen. This was followed by a stimulus that remained on the screen until the participant responded. Participants pressed the ‘Z’ key on the keyboard if they thought the stimulus belonged in Category A, or the ‘M’ key for Category B. In the immediate feedback conditions, participants received feedback after every trial. If the participant correctly categorized the stimulus, they heard a high-pitched tone and then proceeded to the next trial after a 500ms inter-trial interval.

If the participant incorrectly categorized the stimulus, they heard a low pitched tone and proceeded to the next trial after a 4000ms inter-trial interval. Participants were able to take a self-paced break after every 6 trials. In total, participants completed 600 trials.

In the deferred feedback conditions, participants responded to a block of six stimuli, without feedback, with an inter-trial interval of 250ms. At the end of the block of six trials, the feedback was collectively presented. Participants received feedback for correct trials first, hearing a high-pitched tone for every correct categorization response made, each separated by 500ms. A low-pitched tone was then played for every incorrect response made, with each incorrect tone followed by a 4 second ITI. The tones presented were the same as in the immediate feedback condition. The next set of six trials followed after a self-paced break.

### Bayesian analysis

To supplement the classic null-hypothesis significance testing approach, we also calculated Bayes Factors for the ANOVA results we present. This is because, in traditional null-hypothesis significance testing, non-significant results are ambiguous: they could either be due to insufficient statistical power or due to the null hypothesis being correct (Dienes, 2011). It is important to be able to distinguish between these two possibilities.

By convention, if the Bayes Factor is over three then the experiment has found evidence for the experimental hypothesis, whereas if the Bayes Factor is less than a third, the experiment finds evidence for the null hypothesis (Jeffreys, 1961). A Bayes Factor of one indicates that the evidence is exactly neutral with respect to the experimental and null hypotheses (Dienes, 2011). Values between a third and three are typically interpreted as indicating that the experiment was not sensitive enough and no conclusions can be drawn.

The Bayes Factors in this article were calculated according to the procedure recommended by Dienes (2011) using the R script implemented by Baguley and Kaye (2010). The priors were two-tailed normal distributions and a standard deviation of half the mean. In Dienes (2011), the standard deviation of the prior is typically defined as half the mean; this captures the belief that the true mean difference could plausibly take a range of values, but that an effect in the opposite direction to that previously observed is unlikely.

Experiments 1, 2a, and 3a used the means from Smith et al. (2014)’s original study as the prior of our Bayes Factor calculations. In Experiments 2b and 3b, we utilized the means from Experiments 2a and 3a, respectively, as both served to replicate key elements of these earlier studies. All data analyses were conducted in R (R Core Team, 2024).

## Results

We report below the analysis of learning performance after excluding non-learners. There are two ways of excluding non-learners in the COVIS literature. Newell et al. (2010) excluded participants who are not significantly above chance by the final block. In other words, those who scored less than 0.6 on the final 100 trials. In this experiment, this resulted in excluding 11 participants: one from the UD-deferred condition, two from the II- immediate condition and eight from the II-deferred condition. An alternative approach is the method from Smith et al. (2014) where participants were excluded if they performed significantly lower in the last 100 trials compared with the first 100 trials. Unfortunately, Smith et al. did not specify exactly how they calculated this, so here we used a chi-squared test as this seemed most appropriate. In this experiment, this would result in excluding one participant from the UD-immediate condition. Here and in subsequent experiments, we report statistical analyses based on the participants that remain after using the Newell et al. (2010) exclusion criterion. In the Supporting Information, we report the analyses using the Smith et al. exclusion criteria for each experiment reported in this paper. In all cases, the core pattern of results was the same no matter the exclusion criterion used.

Figure 3 shows the results of Experiment 1. As in Smith et al. (2014), the proportion of correct responses in the last 100 trials was compared across conditions in a 2x2 between-subjects ANOVA with the independent variables being category structure (UD/II) and feedback type (immediate/deferred). There was evidence of a significant interaction between category structure and feedback type, *F*(1,71) = 5.53, *p* = .021, *η*_g_^2^ = 0.07, *BF* = 6.52. Welch independent samples t-tests revealed there was a significant drop in accuracy in the deferred feedback condition compared to the immediate feedback condition in the II conditions, *M*_imm_ = 0.82, *SD* = 0.11, *M*_def_ = 0.71, *SD* = 0.06, t(27.4) = 3.74, p <.001, d = 1.24, but not in the UD conditions, *M*_imm_ = 0.85, *SD* = 0.10, *M*_def_ = 0.84, *SD* = 0.08, t(39.2) = .45, p= .65, d = .14. There was a significant main effect of category structure on accuracy, *F*(1,71) = 14.47, *p < .*001, *η*_g_^2^ = 0.17, BF = 351.32, with participants in the UD conditions, *M* = 0.84, *SD* = 0.09, more accurate than II participants, *M* = 0.77, *SD* = 0.10. There was also a statistically significant main effect of feedback type, *F*(1,71) = 8.68, *p* = .004, *η_g_*^2^ = 0.11, BF = 45.67, with participants in deferred feedback conditions performing less accurately, *M* = 0.78, *SD* = 0.10, than participants in the immediate feedback conditions, *M* = 0.84, *SD* = 0.10. Finally, like Smith et al. (2014), we found no significant difference in accuracy for UD and II categorization under immediate feedback, t(35.3) = .92, p = .36, d = .29.

**Fig 3.**
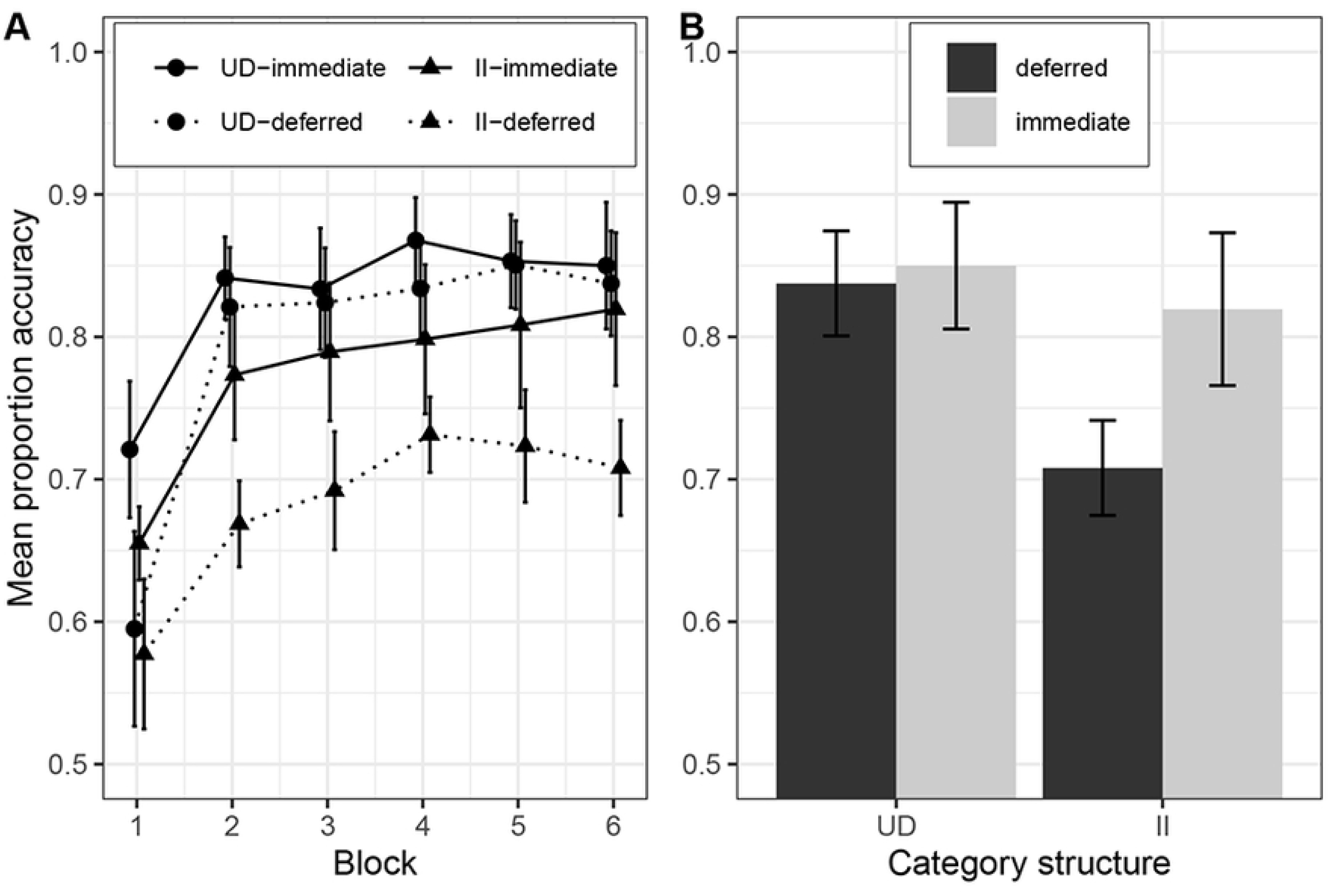
Learning performance in Experiment 1: A) across the experiment, where each block contains 100 trials, and B) in the last block. Category structures: UD = Unidimensional, II = Information-integration.

## Discussion

Experiment 1 successfully replicated the key results from Smith et al. (2014) that were taken as providing strong support for COVIS. There was a significant interaction between category structure and feedback type, with II performance significantly impaired by deferring feedback whereas UD accuracy was statistically unaffected by feedback type. Additionally, unlike Le Pelley et al. (2019) but as in Smith et al. (2014), we found no significant difference in UD and II accuracy in the immediate conditions. The comparable levels of accuracy between the II and UD conditions suggests that the dot stimuli are well controlled in terms of difficulty.

### Experiment 2a

Having successfully replicated Smith et al.’s (2014) finding of a dissociation in the effect that deferred feedback has on UD and II learning, in Experiments 2a and 2b we investigated whether we could replicate Le Pelley et al.’s (2019) key result that for the line stimuli this effect is not present when a CJ category structure is compared to an II structure.

## Method

### Participants and Design

86 University of Exeter students completed the experiment (16 males, mean age = 19.15 years, SD = 2.77). Four participants were excluded due to a data error where their condition was not recoverable. Participants received course credit or £5 remuneration for participation in the study. The study employed a 2 (category structure) X 2 (feedback type) design creating four conditions which participants were randomly allocated to: CJ immediate (n=21); CJ deferred (n=20); II immediate (n=20); and II deferred (n=21).

### Stimuli

As in Le Pelley et al. (2019), we used black line stimuli on a white background varying in length and orientation. We used sets of 600 conjunctive RB and II stimuli so that both dimensions were relevant for learning in the RB and II conditions. These stimuli were generated in the same way as in Le Pelley et al. (the distributions are shown in Table 2) and in Filoteo et al. (2010). The conjunction rule was “short, upright lines belong in category A, and the rest in category B” and the II condition was separated by a diagonal decision boundary. There was a 5% overlap between the categories so that for both the CJ and II tasks the maximum accuracy that could be achieved was 95%.

**Table 2.**
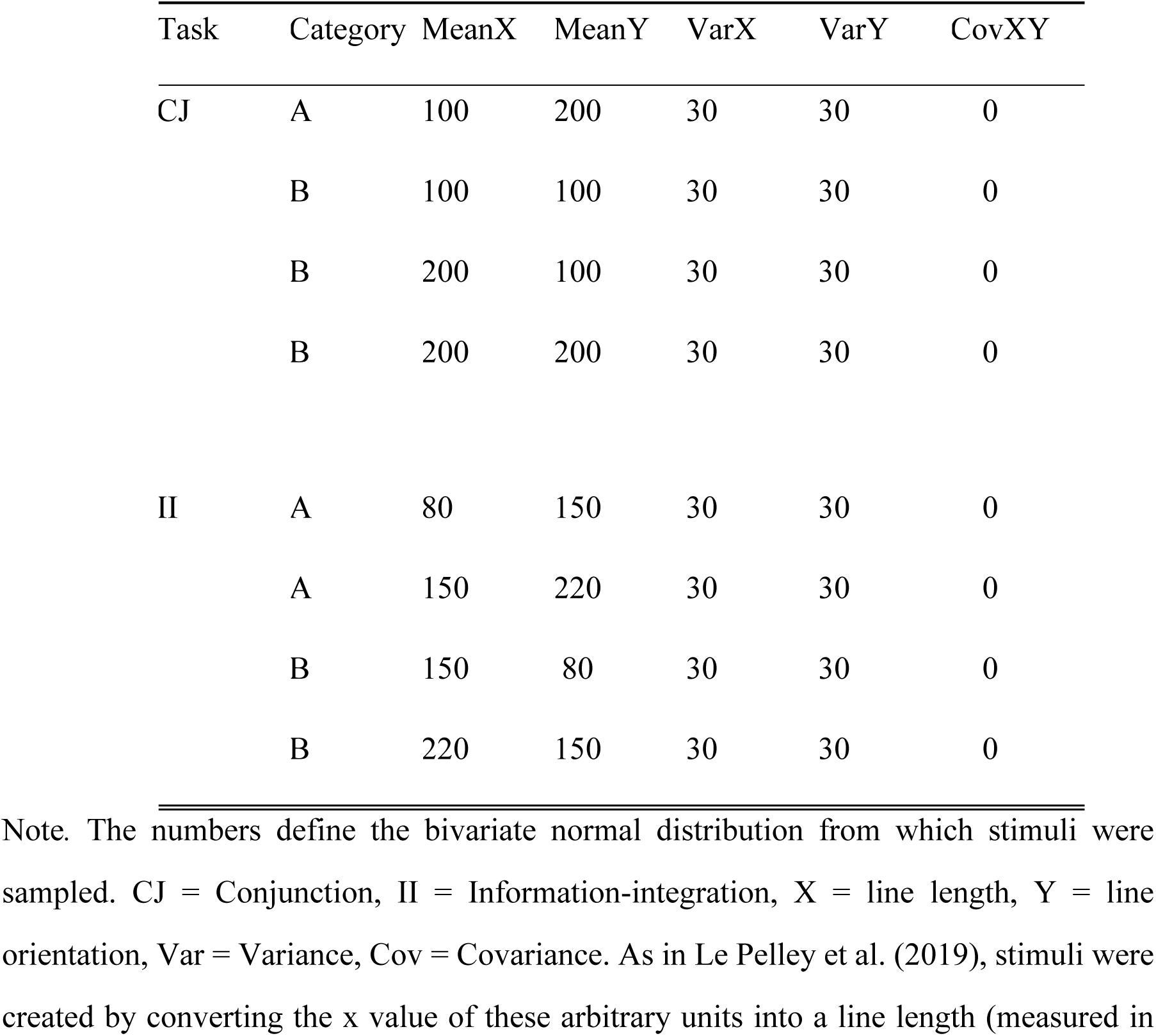
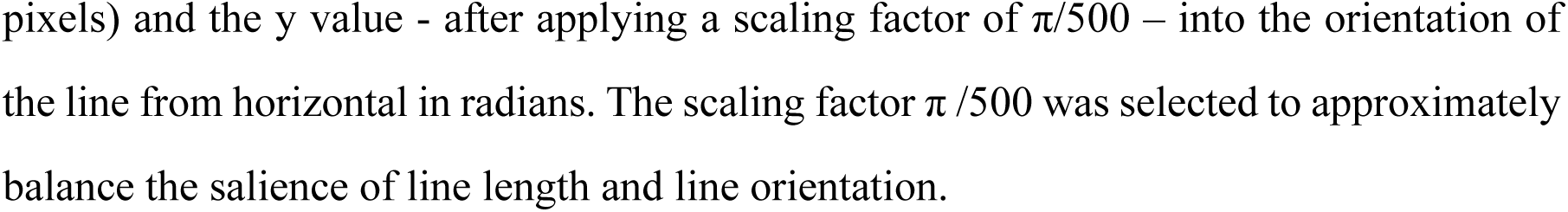
Distributional characteristics of the conjunction RB and II category structures in Experiments 2a and 2b.

| Task | Category | MeanX | MeanY | VarX | VarY | CovXY |
| --- | --- | --- | --- | --- | --- | --- |
| CJ | A | 100 | 200 | 30 | 30 | 0 |
|  | B | 100 | 100 | 30 | 30 | 0 |
|  | B | 200 | 100 | 30 | 30 | 0 |
|  | B | 200 | 200 | 30 | 30 | 0 |
| II | A | 80 | 150 | 30 | 30 | 0 |
|  | A | 150 | 220 | 30 | 30 | 0 |
|  | B | 150 | 80 | 30 | 30 | 0 |
|  | B | 220 | 150 | 30 | 30 | 0 |

Note. The numbers define the bivariate normal distribution from which stimuli were sampled. CJ = Conjunction, II = Information-integration, X = line length, Y = line orientation, Var = Variance, Cov = Covariance. As in Le Pelley et al. (2019), stimuli were created by converting the x value of these arbitrary units into a line length (measured in
pixels) and the y value - after applying a scaling factor of $\pi/500$ – into the orientation of the line from horizontal in radians. The scaling factor $\pi/500$ was selected to approximately balance the salience of line length and line orientation.

### Procedure

The procedure used was identical to Experiment 1.

## Results

We excluded one participant from the CJ-immediate condition, four participants from the CJ-deferred condition, and 7 participants from the II-deferred conditions for failing to meet the learning criterion (Newell et al. 2010). The results of Experiment 2a are shown in Figure 4. We again assessed performance in the last 100 trials across conditions using a 2 x 2 ANOVA with the factors category structure (II/RB) and feedback type (immediate/deferred). We found no significant effect of category structure on accuracy, *F*(1,66) = .77, *p* = .380, *η*_g_^2^ = .01, BF = .10 (CJ, *M* = 0.78, *SD* = 0.08; II, *M* = 0.77, *SD* = 0.09). There was, however, a main effect of feedback type, *F*(1,66) = 40.43, *p < .*001, *η*_g_^2^ = .38, BF = 62.8^7^. with participants in deferred feedback conditions performing less accurately, *M* = 0.71, *SD* = 0.07, than participants in immediate feedback conditions, *M* = 0.82, *SD* = 0.08. However, as in Le Pelley et al., there was no evidence for the key interaction between category structure and feedback type, *F*(1,66) =.24, *p* = .626, *η_g_*^2^ = .004, BF = .14, that we observed in Experiment 1 and Smith et al. (2014) found in their study when comparing the II condition to the UD condition.

**Fig 4.**
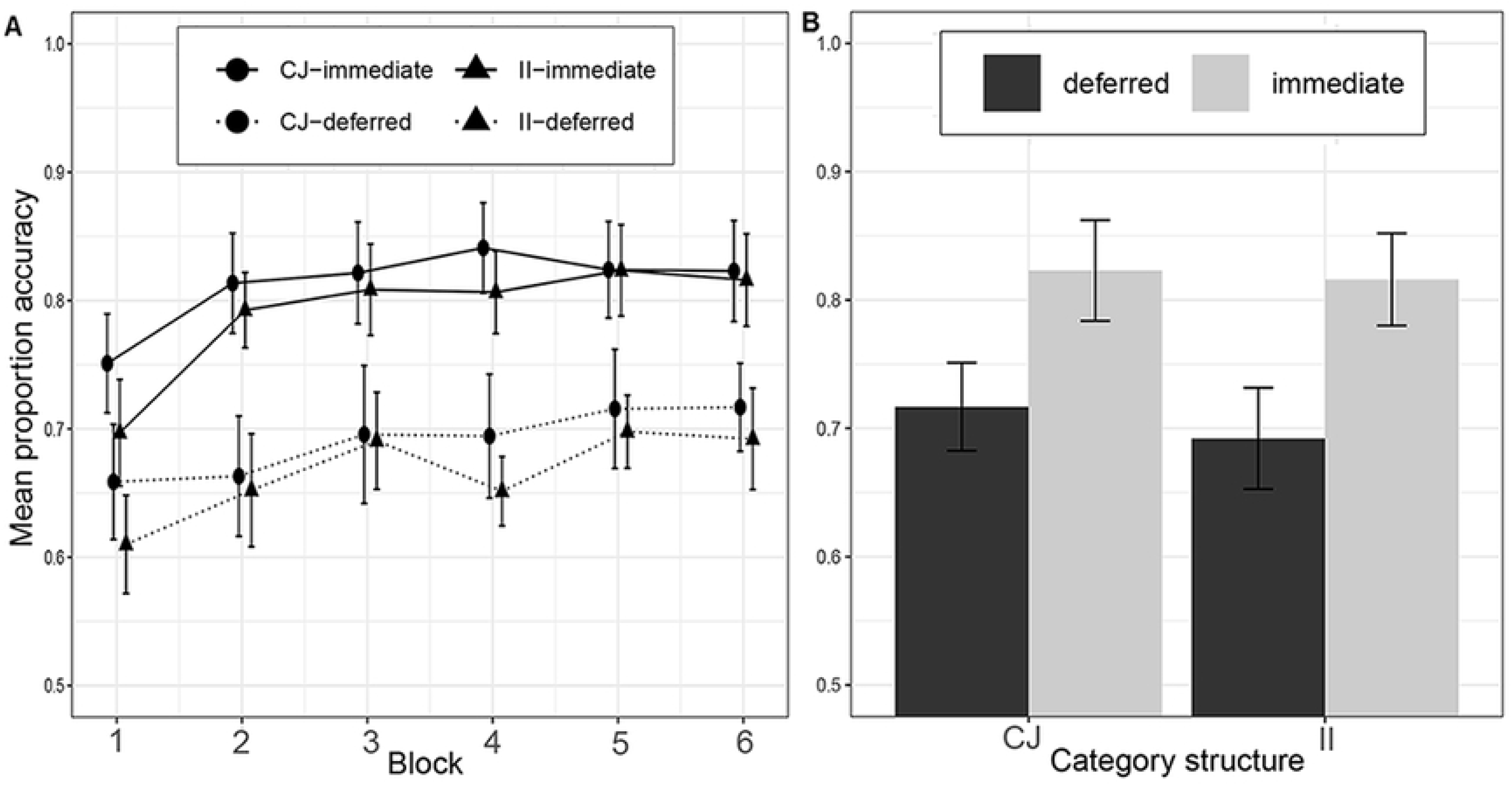
Learning performance in Experiment 2a: A) across the experiment, where each block contains 100 trials, and B) in the last block. Category structures: CJ = Conjunction, II = Information-integration.

## Discussion

This experiment successfully replicated the key result of Le Pelley et al. (2019) by finding that deferred feedback similarly reduced learning performance in both the CJ and II tasks relative to the immediate feedback condition. Our results so far support the idea that the deferred feedback effect is present when the II category structure is compared to a UD structure but not a CJ structure. Nevertheless, given the pivotal role that the CJ condition has in this debate we decided in Experiment 2b to further confirm the result that CJ learning with the line stimuli is impaired by deferring feedback.

## Experiment 2b

### Method

86 University of Exeter students completed the experiment (23 males, mean age = 20.64 years, SD = 4.04). Participants received course credit or £5 remuneration for participation in the study. A between-subjects design was implemented with two conditions: CJ learning with immediate feedback, and CJ learning with deferred feedback. The CJ category structure was the same as in Experiment 2a and we again used the line stimuli. The procedure was identical to the previous experiments.

## Results

We excluded 17 participants from the deferred feedback condition and three participants from the immediate feedback condition for not reaching the learning criterion. The results are displayed in Figure 5. There was a significant main effect of feedback type, *F*(1,63) = 14.56, *η*_g_^2^ = 0.19, *p < .*001, BF = 442, with participants in the immediate condition more accurate, *M* = 0.82, *SD* = 0.08, than those in the deferred feedback condition, *M* = 0.74, *SD* = 0.07.

**Fig 5.**
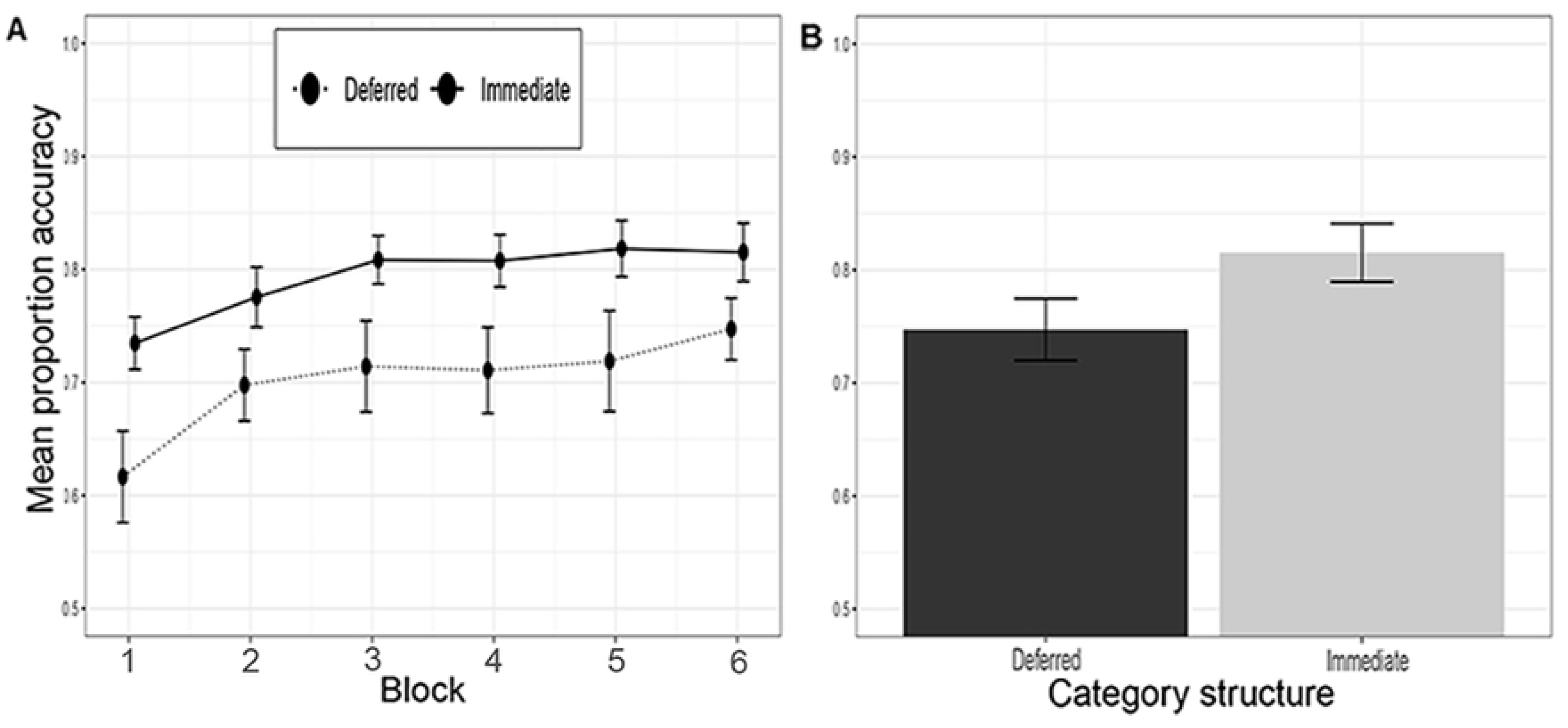
Learning performance in Experiment 2b: A) across the experiment, where each block contains 100 trials, and b) in the last block. Category structures: CJ = Conjunction, II = Information-integration.

## Discussion

In Experiment 2b, we confirmed the finding that participants learning a CJ category structure performed significantly worse when receiving deferred feedback than did participants in the immediate feedback condition. Taken together with the results of Experiment 2a, this indicates that both CJ and II learning can be disrupted by deferred feedback which is consistent with the account put forward by Le Pelley et al. (2019).

### Experiment 3a

In Experiment 1, we replicated the original experiment of Smith et al. (2014) using the dot stimuli. We found a significant interaction as deferring feedback impaired performance on a II task, but left learning of a UD task intact. In Experiments 2a and 2b, we replicated the work of Le Pelley et al. (2019) using the line stimuli. There was no interaction and we found that deferring feedback reduces performance in both II and CJ tasks. In Experiments 3a and 3b, we examine whether our results from Experiments 2a and 2b extend to the dot stimuli used in Smith et al.’s study.

If we again find an absence of an interaction with II and CJ learning being equally impaired by deferred feedback then this would provide considerable support for Le Pelley et al.’s (2019) proposal that Smith et al.’s (2014) original finding was driven by the II task having two relevant dimensions and the UD task only one relevant dimension and that when this confound is removed the effect disappears. On the other hand if the pattern of CJ performance does not generalise to the dot stimuli employed by Smith et al. and II learning is more impaired than CJ learning under deferred feedback then this would be a striking demonstration of the critical influence the type of stimuli have on whether the effect is observed and add a level of complexity to this debate not captured in either the Smith et al. or Le Pelley et al. papers.

## Method

### Participants and Design

84 University of Exeter students completed the experiment. Four participants were excluded due to a data error where their condition was not recoverable. Participants received course credit or £5 remuneration for participation in the study. The study employed a 2 (category structure) X 2 (feedback type) design with participants randomly allocated to one of the four conditions: CJ immediate (n=21); CJ deferred (n=21); II immediate (n=22); and II deferred (n=20).

### Stimuli

The stimuli were the dot stimuli used by Smith et al. (2014) in their original experiment and in our Experiment 1. As in previous experiments there were 600 unique stimuli in the II and CJ conditions. The stimuli for each participant were randomly generated using the procedure outlined in Experiment 1 (the distributions are displayed in Table 3).

**Table 3.**
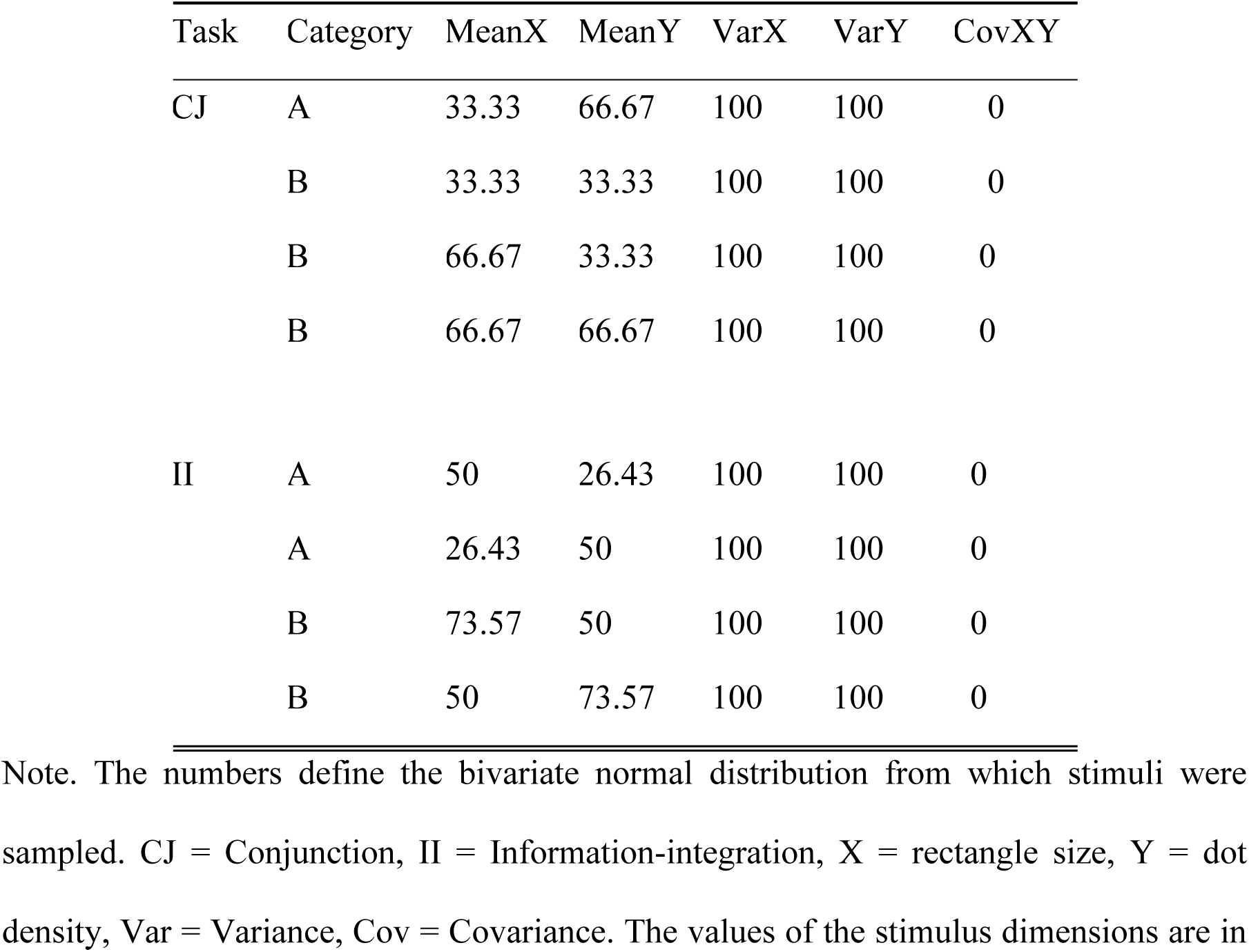
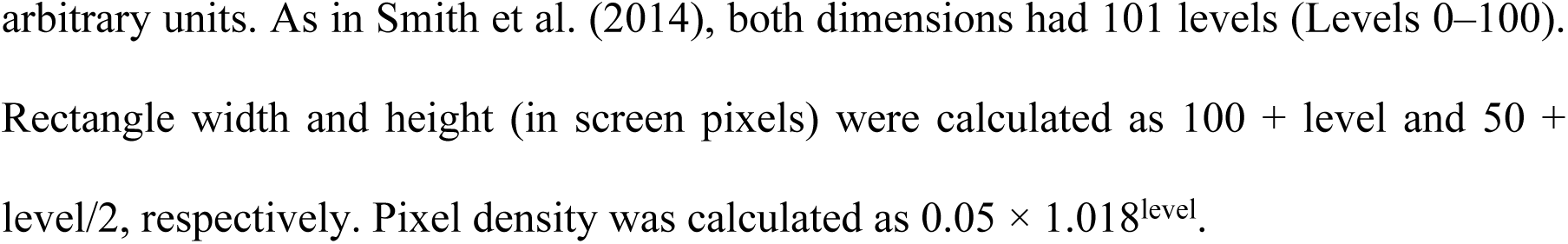
Distributional characteristics of the conjunction RB and II category structures in Experiments 3a and 3b.

| Task | Category | MeanX | MeanY | VarX | VarY | CovXY |
| --- | --- | --- | --- | --- | --- | --- |
| CJ | A | 33.33 | 66.67 | 100 | 100 | 0 |
|  | B | 33.33 | 33.33 | 100 | 100 | 0 |
|  | B | 66.67 | 33.33 | 100 | 100 | 0 |
|  | B | 66.67 | 66.67 | 100 | 100 | 0 |
| II | A | 50 | 26.43 | 100 | 100 | 0 |
|  | A | 26.43 | 50 | 100 | 100 | 0 |
|  | B | 73.57 | 50 | 100 | 100 | 0 |
|  | B | 50 | 73.57 | 100 | 100 | 0 |

Note. The numbers define the bivariate normal distribution from which stimuli were sampled. CJ = Conjunction, II = Information-integration, X = rectangle size, Y = dot density, Var = Variance, Cov = Covariance. The values of the stimulus dimensions are in
arbitrary units. As in Smith et al. (2014), both dimensions had 101 levels (Levels 0–100). Rectangle width and height (in screen pixels) were calculated as $100 + \text{level}$ and $50 + \text{level}/2$ , respectively. Pixel density was calculated as $0.05 \times 1.018^{\text{level}}$ .

### Procedure

The procedure was identical to the preceding experiments.

## Results

One participant was excluded from the CJ-immediate condition, two participants from the CJ-deferred condition, four participants from the II-immediate condition and 11 participants from the II-deferred condition for failing to meet the learning criterion. Figure 6 show the results of Experiment 3a. There was a significant interaction between feedback type and category structure, *F*(1,65) = 7.15, *p* = .009, *η*_g_^2^ = 0.10, BF = 14.17. Welch independent samples t-tests revealed that deferred feedback reduced accuracy in II category learning relative to immediate feedback, *M*_imm_ = 0.84, *SD* = 0.07, *M*_def_ = 0.70, *SD* = 0.09, t(15.4) = 4.37, p <.001, d = 1.83, but there was no significant effect of feedback on CJ category learning, *M*_imm_ = 0.76, *SD* = 0.09, *M*_def_ = 0.72, *SD* = 0.06, t(35.3) = 1.55, p = .129, d = .48. There was a significant main effect of feedback condition, *F*(1,65) = 21.09, *p < .*001, *η*_g_^2^ = 0.24, BF = 13025, with participants learning better with immediate feedback, *M* = 0.80, *SD* = 0.09, than deferred feedback, *M* = 0.71, *SD* = 0.07. The main effect of category structure was not statistically significant, *F*(1,65) = 1.98, *p* = .164, *η*_g_^2^ = 0.03. BF = .05 (II = .79, SD = .10; CJ = .74, SD = .08).

**Fig 6.**
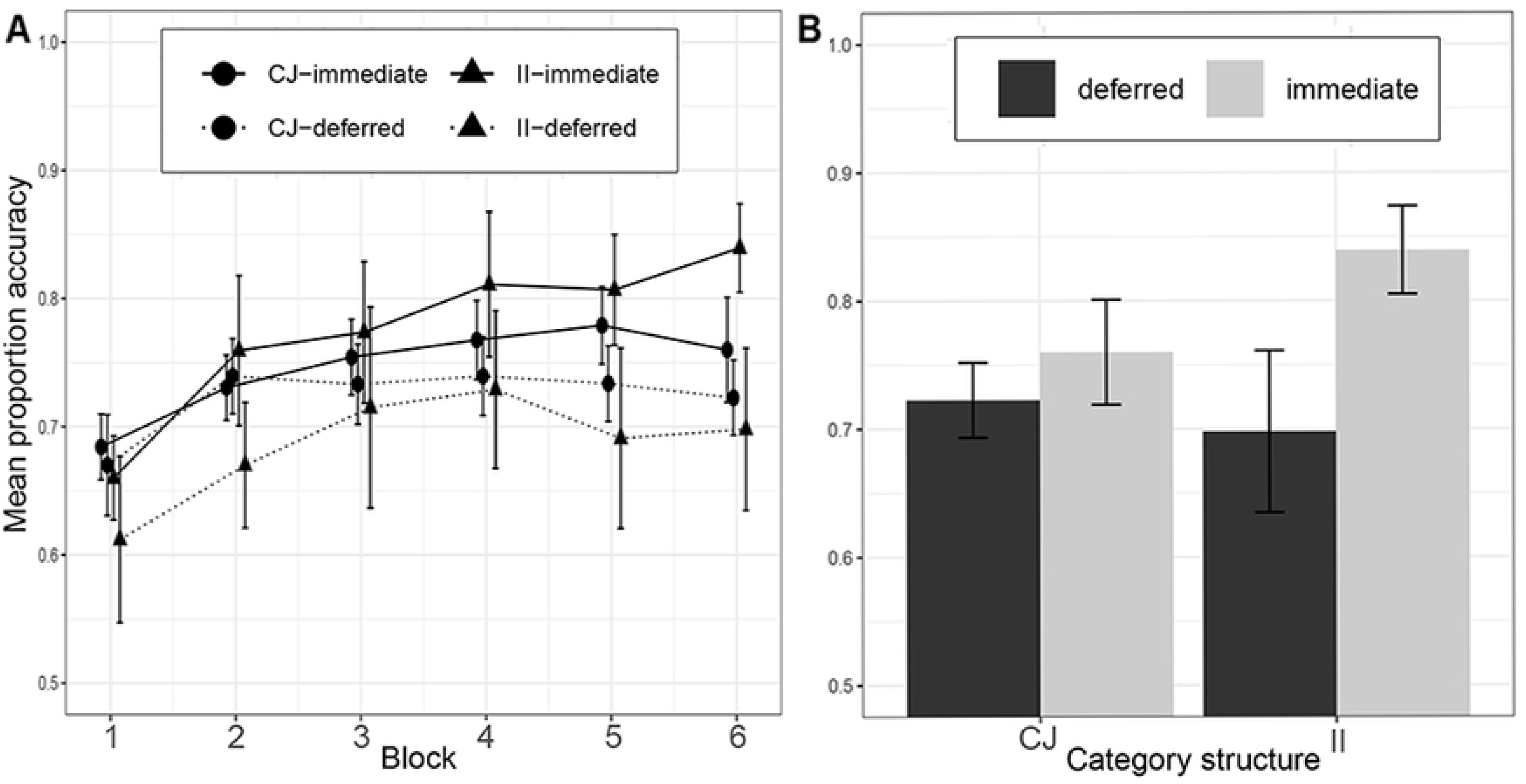
Learning performance in Experiment 3a: A) across the experiment, where each block contains 100 trials, and b) in the last block. Category structures: CJ = Conjunction, II = Information-integration.

## Discussion

In this experiment, we found that deferred feedback impaired II category learning to a significantly greater extent than CJ learning. This pattern of results is different from what we observed with the line stimuli in Experiments 2a and 2b and what Le Pelley et al. (2019) found in their paper and suggests that the type of stimuli used appears to be driving the discrepancy in results between the two studies rather than whether the II structure is compared to a UD or a CJ task. In contrast, these results appear consistent with COVIS by demonstrating that the effect of deferred feedback is greater for II than RB category structures. Nevertheless, before drawing any strong inferences about these results we ran a direct replication of this study to see whether this result is reliable.

## Experiment 3b

### Method

#### Participants and Design

84 University of Plymouth students completed the experiment. The study employed a 2 (category structure) X 2 (feedback type) design creating four conditions: CJ immediate (n=24); CJ deferred (n=25); II immediate (n=23); and II deferred (n=24). This experiment received ethical approval from the University of Plymouth, Psychology Ethics Committee.

#### Stimuli and Procedure

The stimuli were generated in the same way as in Experiment 3a and the procedure was identical to all the previous experiments.

## Results

Two participants from the CJ-immediate condition, six participants from the CJ-deferred condition, and three participants from the II-deferred condition were excluded for being non-learners. The results of Experiment 3b are shown in Figure 7. As in Experiment 3a, there was a significant interaction between feedback type and category structure, *F*(1,81) = 7.27, *p* = .009, *η*_g_^2^ = 0.08, BF = 18.01, indicating that the impairment of deferred feedback on II category learning, *M*_imm_ = 0.82, *SD* = 0.07, *M*_def_ = 0.70, *SD* = 0.07, t (42.0 = 5.26, p<.001, d 1.58) was greater than on CJ category learning, *M*_imm_ = 0.75, *SD* = 0.06, *M*_def_ = 0.72, *SD* = 0.06, t (38.8) = 1.77, p = .08, d = .55. There was also a significant effect of feedback condition, *F*(1,81) = 25.44, *p < .*001, *η*_g_^2^ = 0.24, BF = 97826, with participants learning better with immediate feedback (M = .79, SD = .08) than deferred feedback (M = .71, SD = .06). There was no significant main effect of category structure, *F*(1,81) = 3.03, *p* = .086, *η*_g_^2^ = 0.04, BF = 3.11 (CJ, M = .74, SD = .06; II, M= .76, SD = .09).

**Fig 7.**
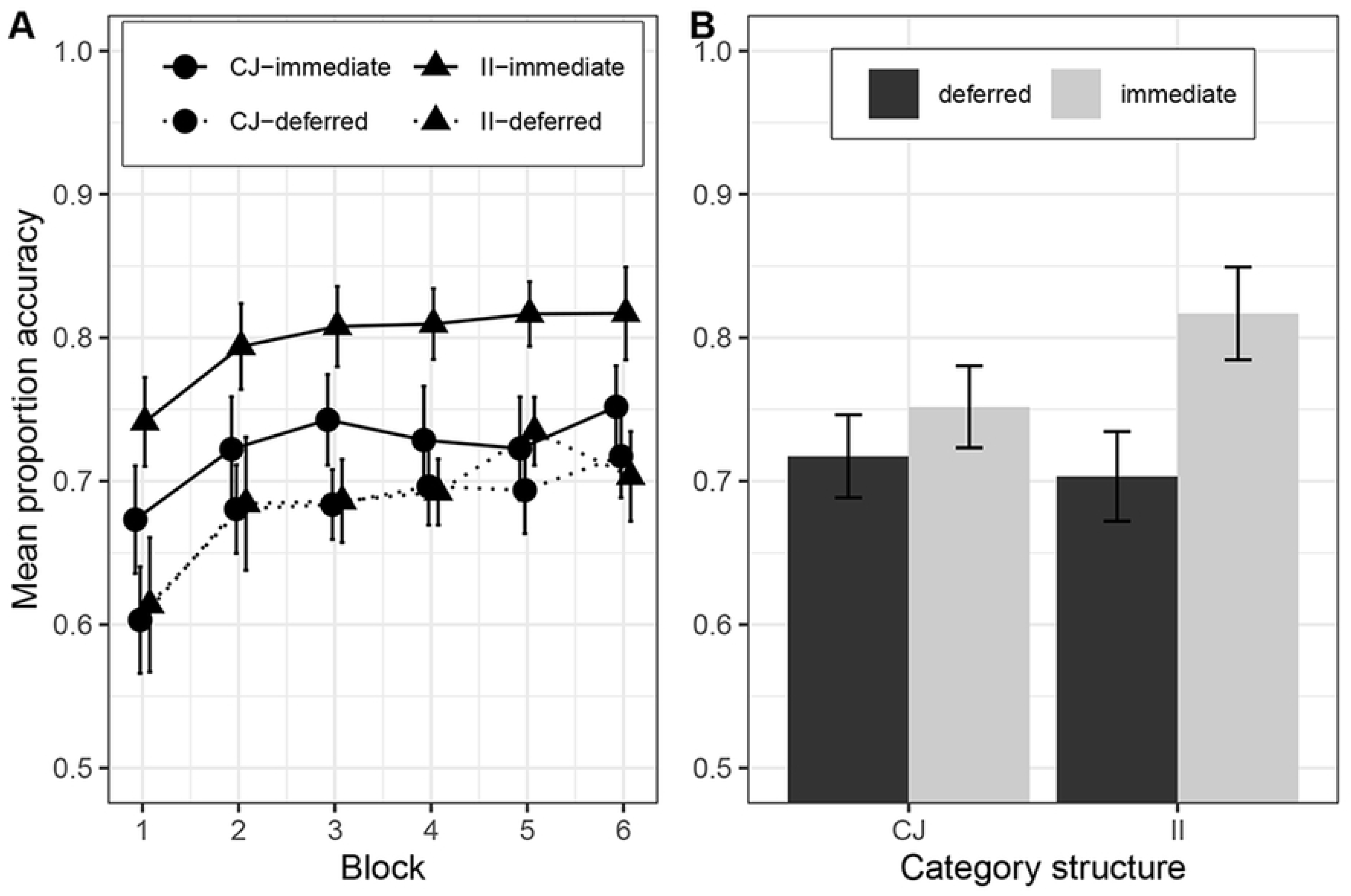
Learning performance in Experiment 3b: A) across the experiment, where each block contains 100 trials, and b) in the last block. Error bars are 95% confidence intervals. Category structures: CJ = Conjunction, II = Information-integration.

## Discussion

Experiment 3b successfully replicated the results of Experiment 3a by again finding with the dot stimuli a significant interaction between category structure and feedback type – deferred feedback impaired learning of an II task but not a CJ task. These findings appear, at least on the surface, to challenge Le Pelley et al.’s (2019) idea that the effect of deferred feedback on category learning observed in Experiment 1 and in Smith et al. (2014) was driven by the greater number of relevant dimensions in the II task than the UD task because here both the rule-based and information-integration tasks had two relevant dimensions and yet we have obtained the same dissociation of deferred feedback on II and RB category structures as Smith et al.

## General discussion

This paper revisited Smith et al.’s (2014) finding that deferred feedback significantly impaired learning of a two-dimensional II category structure but had no adverse effect on a UD RB category structure. Smith et al. took this result as providing some of the strongest evidence to date in support of the COVIS dual-process categorization model which proposes that there are separate implicit and explicit category learning systems. They argued that deferred feedback selectively impaired II categorization, as this is thought to engage the implicit learning system which requires prompt feedback, but left UD rule-based categorization intact, because this task is believed to recruit the explicit system which does not need immediate feedback. This claim has subsequently been challenged by Le Pelley et al. (2019) who argued that this effect was driven by the II task being more cognitively demanding than the UD task, as whilst only one dimension was relevant for learning the UD task, the II task had two relevant dimensions. This account argues that deferred feedback simply impairs the more difficult task to a greater extent than the easier task. Le Pelley et al. provided support for this by demonstrating that deferred feedback impaired CJ RB learning and II learning, which are matched for the number of relevant dimensions, to the same extent.

In Experiments 1 and 2 we provided the first direct replication of the results of Smith et al. and Le Pelley et al. and confirmed that their results were reliable. However, one notable aspect of Le Pelley et al.’s study was that the stimuli they used – line stimuli varying in length and orientation – were very different from the “dot stimuli” that Smith et al. used – rectangles varying in size and the density of dots – raising the possibility that the reason for the discrepancy in findings between the two studies was not due to whether category structures were matched for the number of relevant dimensions but due to the characteristics of the stimuli used. To discriminate between these two possibilities, in Experiment 3 we used the same CJ and II structures as Le Pelley et al. but now with the dot stimuli that Smith et al. had used. With this approach, consistent with Smith et al., we found a significant interaction with deferred feedback impairing II learning to a greater extent than CJ learning.

This result might seem surprising given that previous evidence supports Le Pelley et al.’s (2019) contention that COVID related effects can result from failing to match the number of dimensions across II and rule-based tasks (e.g., Carpenter et al., 2016; Edmunds et al., 2015) so there were good reasons for thinking that the same explanation might apply to Smith et al.’s (2014) results particularly given Le Pelley et al.’s own findings. However, the results of our Experiment 3 indicate that the difficulty account cannot straightforwardly provide a complete explanation for Smith et al.’s results because the deferred feedback effect was present with the dot stimuli when the number of relevant dimensions was matched across the II and RB conditions. Instead, our results highlight the key role that the type of stimuli employed can have on whether Smith et al.’s deferred feedback effect is obtained.

In the COVIS literature, significant attention has been given to how different types of category structures influence the pattern of behavioral results. However, there has been little exploration of whether the type of stimuli might similarly affect these outcomes. We suspect part of the reason for this is that most COVIS-related experiments use very similar and simple stimuli such as the line stimuli we have used here or Gabor patches varying on bar width and orientation. Indeed, outside Smith et al.’s study, the dot stimuli have seldom been used in COVIS-related work or indeed categorization research more generally. There are valid reasons for focusing on the same type of highly controlled stimuli, but this work raises the question of whether results from such stimuli will generalize to other types, even when using the same category structures. Our findings suggest that while replicating results within the same stimulus set is important, greater emphasis should be placed on ensuring that effects also transfer to other stimulus sets in order to better establish their generality.

Whilst we consider the primary contribution of our paper is highlighting the key role that the type of stimuli play in whether the deferred feedback effect is observed, it is worth considering the implication of our results to the multiple category learning systems debate. Dual process theorists would likely see our results as being consistent with the idea that there are separate explicit and implicit category learning systems. However, the deferred feedback effect occurs in more limited conditions than those described in Smith et al.’s paper. Furthermore, COVIS currently does not appear to consider stimulus-specific effects so it may need some revision to explain why the deferred feedback effect emerges for the dot stimuli but not the line stimuli.

On the other hand, single process theorists would likely question how compelling this evidence is for dissociable category learning systems by raising concerns about the dot stimuli. One notable aspect of the dot stimuli is that they appear much more perceptually complex than the line stimuli and indeed other stimuli commonly used in COVIS research. There is, of course, nothing wrong with the use of perceptually complex stimuli and one can certainly argue, as we have done above, that it is informative to use a rich mix of stimuli but equally whilst the line stimuli have a straightforward physical-to-psychological mapping, the properties of the novel dot stimuli are not well understood given how little they have been used in past research. Le Pelley et al. (2019) raised the concern that the dot stimuli may be susceptible to an emergent dimension. Similarly, it may just be harder to detect precisely what the relevant dimensions are for the perceptually complex dot stimuli. In either case this may encourage people to use what would be a suboptimal UD strategy for the CJ structure. If participants are favouring a UD strategy for the CJ structure then this would provide an explanation for why the CJ/dot stimuli show the same pattern of results as the UD/dot stimuli under deferred feedback but the CJ/line stimuli, where using both dimensions might be an easier strategy to apply, do not. In many previous COVIS- related studies, decision bound analysis has been conducted to provide insight into categorization strategies that participants are using. However, in recent years research has highlighted the poor validity of decision bound analysis in this regard (e.g., Donkin et al., 2015; Edmunds et al., 2018). As a result, we chose not to conduct strategy analyses, given the high likelihood of misclassifying participants’ strategies and then drawing incorrect inferences. One limitation of this explanation, though, is if the dot stimuli encourage a UD strategy for the CJ structure then by the same reasoning they might be expected to encourage a UD strategy for the II structure so it remains unclear from the difficulty account why deferred feedback would more markedly affect II category learning. Consequently, the results presented here appear to pose a challenge to the single system account. To gain a clearer understanding of how strongly this effect supports COVIS, future research should focus on a more detailed characterization of the dot stimuli and investigate the deferred feedback effect using additional stimulus sets.

In summary, the current paper provides a striking example of how the type of stimuli can have a critical influence on whether a behavioral effect is observed in a category learning task. We found that deferred feedback impaired II learning to a greater extent than CJ learning with one set of stimuli but this effect disappeared when a different set of stimuli were used. Previous COVIS-related work has focused on the differential impact that different category structures have on categorization performance under various manipulations. Whilst this work does not challenge the view that this is an important consideration, it does highlight that the nature of the stimuli employed in COVIS research needs more attention. Our results suggest that to avoid effects that are only valid under specific conditions, future research should ensure that effects are tested across a variety of stimuli to establish their generality and increase confidence in their robustness.

## Acknowledgements

The work was supported by a South West Doctoral Training Centre (SWDTC) Economic and Social Research Council (ESRC) Studentship Award (ES/J50015X/1) to the second author.

## Notes

### Competing Interest Statement

The authors have declared no competing interest.

